## Supplementary Information for "Profiling of the *Helicobacter pylori* redox switch HP1021 regulon using a multi-omics approach"

**This PDF file includes:**

**Supplementary Material and methods**

**Supplementary Table 4 to 5**

- Tab. S4. Strains, plasmids and protein used in this study.
- Tab. S4. Primers used in this study.

**Supplementary Figure 1 to 12**

- Fig. S1: Identification of HP1021 binding sites on *H. pylori* N6 genome by ChIP-seq.
- Fig. S2: HP1021 controls *vacA* expression.
- Fig. S3: The reproducibility of biological replicates in omics data.
- Fig. S4: Overview of the gene regulation mediated by HP1021 revealed by RNA-seq.
- Fig. S5: Overview of the protein level regulation mediated by HP1021 revealed by MS/LC-MS.
- Fig. S6: *H. pylori* N6 Clusters of Orthologous Groups (COG).
- Fig. S7: HP1021 controls *katA* and *kapA* expression.
- Fig. S8: HP1021 controls *rocF* expression.
- Fig. S9: Analysis of DNA uptake by *H. pylori* P12.
- Fig. S10: HP1021 controls tRNA expression.
- Fig. S11: HP1021 controlled glucose uptake via GluP transporter.
- Fig. S12: The model of glucose metabolism in *H. pylori* N6.

**Supplementary References**

**Supplementary Material and methods**

**Electrophoretic mobility shift assay (EMSA)**

PCR amplified DNA probes in two steps. DNA fragments were amplified in the first step using unlabelled primers P29-P30 (p*comB8*) or P31-P32 (p*vacA*) and a *H. pylori* N6 genomic DNA template (Table S3 and S4). The forward primers were designed with overhangs complementary to P33, FAM–labelled primer. The unlabelled fragment was purified and used as a template in the second round of PCR using the P33 FAM-labelled primer and the reverse primer P30 or P32 used in the first step. The *oriC2* region was amplified by PCR using P33-P34 primer pairs and a specific pori2 template (Table S3), giving the FAM-*oriC2* probe. FAM-labelled DNA (5 nM) was incubated with the HP1021 protein at 37 °C for 20 min in Tris buffer (50 mM Tris–HCl, pH 8.0; 100 mM NaCl and 0.2% Triton X-100). The complexes were separated by electrophoresis on a 4% polyacrylamide gel in 0.5 × TBE (1 × TBE: 89 mM Tris, 89 mM borate and 2 mM EDTA) at 10 V/cm in the cold room (approx. 10 °C). The gels were analyzed by a Typhoon 9500 FLA Imager and ImageQuant software.

**Catalase activity**

*H. pylori* P12 liquid cultures at the late logarithmic phase of growth ($OD_{600}$ = 0.8-1.0) were collected by centrifugation (5 min, 4,000 × g) to obtain $OD_{600}$ = 1 (approx. $2 \times 10^8$ cells/ml) per sample and resuspended in 1 ml of 1 × PBS. 10 µl of cells' suspension of each strain were simultaneously mixed with 10 µl of 30% $H_2O_2$. The catalase activity was measured by observation of the production of air bubbles and captured by a camera.

**Construction of the *H. pylori* mutant strains**

*H. pylori* mutant strains were constructed using a homologous recombination approach as described by Ge and Taylor[1]. Briefly, *H. pylori* cells were plated from the stock to CBA-B plates and incubated under microaerobic conditions for 24 h. Next, cells were plated on CBA-B plates and incubated for approximately 5 h in the form of a micro-lawn of approx. 10 mm in diameter. Then, the micro-lawn was spotted with 10 µg of purified recombinant plasmids and grown under microaerobic conditions. After 24 h of growth, the culture was plated on a selective medium and grown for five days to obtain single colonies of transformants.

*H. pylori* P12 ΔHP1021

The *H. pylori* P12 ΔHP1021 mutant in which the HP1021 gene was deleted from the chromosome was constructed as described previously for the N6 strain using the pTZ57R/TΔHP1021 plasmid[2]. The allelic exchange was verified by PCR using the P13-P14 primers homologous to the chromosomal regions external to the designed recombination sites. The lack of HP1021 was verified by Western blot analysis using a rabbit polyclonal anti-HP1021 antibody[2] (Fig. S7c-d).

*H. pylori* P12 COM/HP1021

The *H. pylori* P12 COM/HP1021 mutant with the restored HP1021 gene on the chromosome was constructed as described previously for the N6 strain using the pUC18/HP1021com plasmid[2]. The allelic exchange was verified by PCR using the P13-P14 primers homologous to the chromosomal regions external to the designed recombination sites. Additionally, the presence of HP1021 was verified by Western blot analysis using a rabbit polyclonal anti-HP1021 antibody[2] (Fig. S7c-d).

*H. pylori* N6 Δ*gluP*

The *H. pylori gluP* deletion construct (pUC18/ΔgluP) was prepared as follows (Fig. S11c). The upstream and downstream regions of *gluP* were amplified by PCR using the P1-P2 and P5-P6 primer pairs, respectively and *H. pylori* 26695 genomic DNA as a template. The *aphA-3* cassette was amplified using the P3-P4 primer pair; pTZ57R/TΔHP1021[3] was used as a template. The resulting fragments were purified on an agarose gel. Subsequently, the PCR-amplified fragments and the SmaI digested vector pUC18 were ligated according to the method described by Gibson in 2009[4]. *E. coli* DH5α competent cells were transformed by heat shock. The DNA fragment cloned in pUC18 was sequenced. Subsequently, *H. pylori* N6 was transformed with the pUC18/ΔgluP plasmid, and the transformants were selected by plating on CBA-B plates supplemented with kanamycin. The allelic exchange was verified by PCR using the P10-P11 primer pair homologous to the chromosomal regions external to the designed recombination sites. The lack of *gluP* was verified by PCR using the P7-P12 primer pair (Fig. S11e).

*H. pylori* N6 COM/*gluP*

The *H. pylori gluP* complementation strain was constructed as follows (Fig. S11c). The region upstream of *gluP* and the *gluP* gene was amplified by PCR using the P1-P7 primer pair, while the downstream region flanking *gluP* was amplified by PCR using the P5-P6 primer pair; both regions were amplified using *H. pylori* 26695 genomic DNA as a template. The *cat* cassette was amplified using the P8-P9 primer pair; pUC18/HP1021com[2] was used as a template. The resulting fragments were purified on an agarose gel. Subsequently, the PCR-amplified fragments and the SmaI digested vector pUC18 were ligated according to the method described by Gibson in 2009[4]. *E. coli* DH5α competent cells were transformed by heat shock. The cloned insert was sequenced. Subsequently, the obtained plasmid was used in the natural transformation of *H. pylori* N6. The transformants were selected by plating on CBA-B plates supplemented with chloramphenicol. The allelic exchange was verified by PCR using the P10-P11 primers homologous to the chromosomal regions external to the designed recombination sites. Additionally, the presence of *gluP* was verified by PCR using the P7-P12 primer pair (Fig. S11e).

**Supplementary Table**

**Table S4. Strains, plasmids and protein used in this study.**

| Strain | Relevant features | Reference/source |
|---|---|---|
| *E. coli* DH5α | *supE44*, *hsdR17*, *recA1*, *endA1*, *gyrA1*, *gyrA96*, *thi-1*, *relA1* | [5] |
| *E. coli* BL21 | F-, *ompT*, *hsdS (rB-*, *mB-)*, *gal*, *dcm* | GE Healthcare |
| *H. pylori* 26695 | Parental strain | [6] |
| *H. pylori* N6 | Parental strain | [7] |
| *H. pylori* N6 ΔHP1021 | Δ*HP1021*::*aphA-3*; N6 with *HP1021* exchanged to *aphA-3* cassette | [2] |
| *H. pylori* N6 COM/HP1021 | (Δ*HP1021*::*aphA-3*):Δ*HP1021-cat*); N6 Δ*HP1021* in which *aphA-3* was exchanged to *HP1021* and *cat* cassette | [2] |
| *H. pylori* P12 ΔHP1021 | Δ*HP1021*::*aphA-3*; N6 with *HP1021* exchanged to *aphA-3* cassette | This study |
| *H. pylori* P12 COM/HP1021 | (Δ*HP1021*::*aphA-3*):Δ*HP1021-cat*); N6 Δ*HP1021* in which *aphA-3* was exchanged to *HP1021* and *cat* cassette | This study |
| *H. pylori* N6 ΔgluP | Δ*gluP*::*aphA-3*; N6 whit *gluP* exchanged to *aphA-3* cassette | This study |
| *H. pylori* N6 COM/gluP | (Δ*gluP*::*aphA-3*):Δ*gluP-cat*); N6 Δ*gluP* in which *aphA-3* was exchanged to *HPgluP* and *cat* cassette | This study |

| Plasmid | Relevant features | Reference/source |
|---|---|---|
| pUC18 | Cloning vector, $Amp^R$ | Thermo Fisher Scientific |
| pUC18/ΔgluP | pUC18 derivative containing *gluP* flanking regions for allelic exchange of *gluP* for *aphA-3* | This study |
| pUC18/ COM/gluP | pUC18 derivative containing *gluP* flanking regions and *gluP* for allelic exchange of *aphA-3* for *gluP-cat* | This study |
| pTZ57R/TΔHP1021 | pTZ57R/T derivative containing *HP1021* flanking regions for allelic exchange of *HP1021* for *aphA-3* | [3] |
| pUC18/COM/HP1021 | pUC18 derivative containing *HP1021* flanking regions and *HP1021* for allelic exchange of *aphA-3* for *HP1021-cat* | [2] |
| pET28/StrepHP1021 | pET28Strep derivative containing the *HP1021* gene amplified with primers P-3/P-4 and cloned between BamHI and XhoI sites | [2] |
| pori2 | A pOC170 derivative containing *oriC2* | [8] |

| Recombinant protein | Relevant features | Reference/source |
|---|---|---|
| StrepHP1021 | Recombinant, *H. pylori* HP1021 protein, Strep-tagged at N-terminus, purified from *E. coli* | [2] |

**Table S5. Primers used in this study.**

| Oligo name | Sequence (5' → 3') |
|---|---|
| **P1** | gtcgactctagaggatccccgtaaaggtgaagtgccttatc |
| **P2** | gggtataagcaagaagaaaac |
| **P3** | gttttcttcttgcttatacccgctagcctaaaacaattcatccagtaaaata |
| **P4** | ctcaataaaaaaggagaaggtttaattaaggatcctgactaactaggagga |
| **P5** | accttctcctttttattgag |
| **P6** | cgaattcgagctcggtacccgcaatgcgatttatccggtg |
| **P7** | ttaggagttttcttcttgcttatac |
| **P8** | agcaagaagaaaactcctaacggaatttacggaggataaa |
| **P9** | gttttcttcttgcttatacccttacgccccgccctgccact |
| **P10** | atgcaaaaaacttctaacactctg |
| **P11** | ttaacgctcttttgcttgcc |
| **P12** | tccccaagcaaagtcgtgag |
| **P13** | gctcaccacgagcggcgatttg |
| **P14** | gctttaaaatcttcatcaaattgg |
| **P15** | ctagcggattctctcaatgtcaa |
| **P16** | ggagtacggtcgcaagattaaa |
| **P17** | tcaagtcttcaaagcgttgccacac |
| **P18** | aatgaagcgttggctcgttcgctc |
| **P19** | gcctttgacccaataatgcc |
| **P20** | ccaataaaaccccagataaaccc |
| **P21** | gttttctaaatgcttttcttggtgg |
| **P22** | gggacttgtgtttttggttg |
| **P23** | tcagtcgacgctctttttaaggggctttg |
| **P24** | tttttgcataccttctccttttt |
| **P25** | gtgatgctgaccaatgctcc |
| **P26** | tttctagtctaaagtcgcaccc |
| **P27** | catcgcgcaaaccatctcgc |
| **P28** | gacggacaataaggggcgct |
| **P29** | ggagtaagaatagcttcgaatcgaaagcgtctctaaagaaaa |
| **P30** | cgggatcctcatgcgagatttaacctgt |
| **P31** | ggagtaagaatagcttcgaatctatatatttatagccttaatcg |
| **P32** | cggcttgtttgagcccccag |
| **P33** | FAM-ggagtaagaatagcttcgaat* |
| **P34** | catcgataggatatcctggg |

* - FAM, 6-fluorescein amidite.

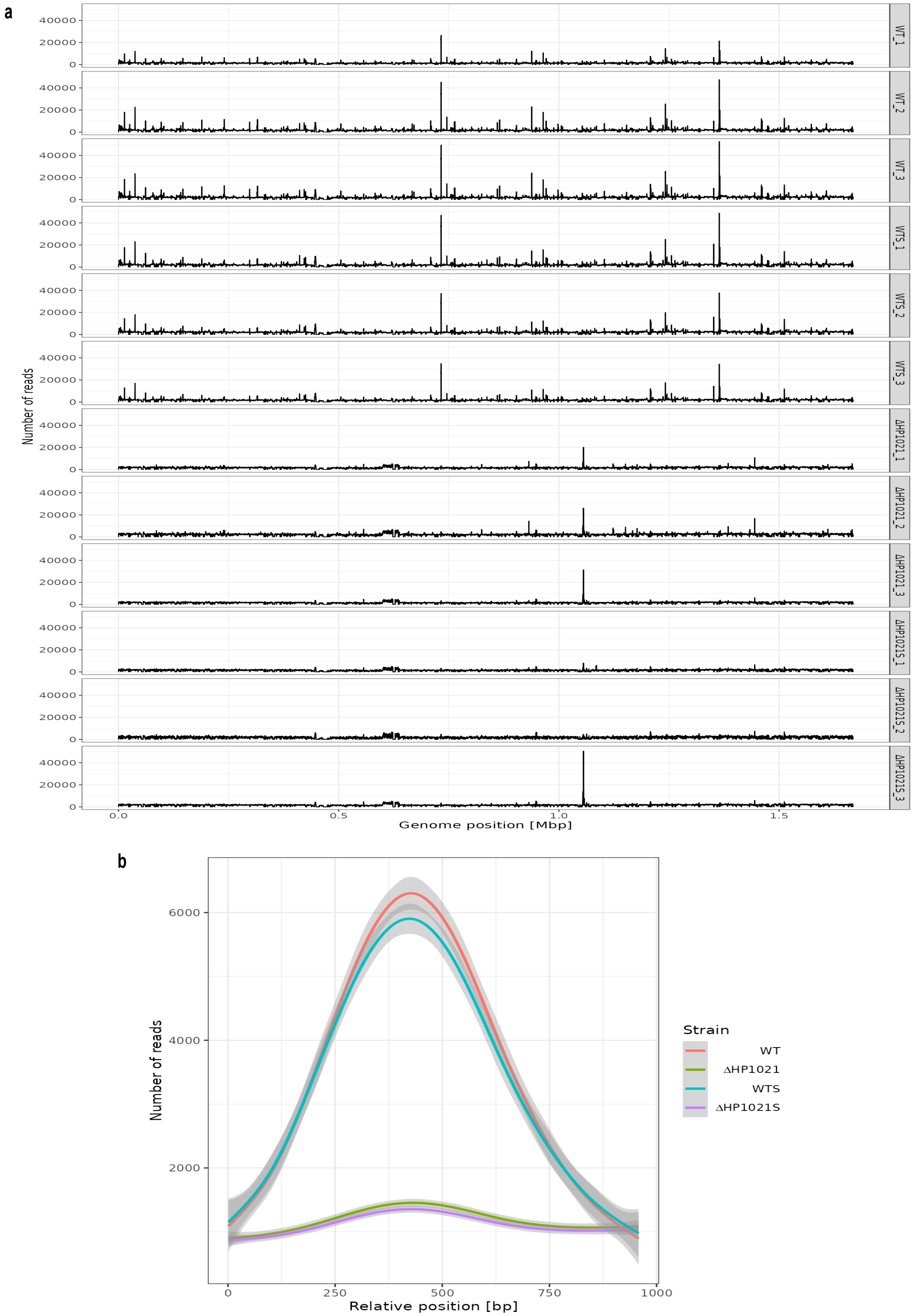


**Fig. S1: Identification of HP1021 binding sites on *H. pylori* N6 genome by ChIP-seq. a** ChIP-seq analysis of HP1021 binding to *H. pylori* N6 chromosome. The ChIP-seq reads for HP1021 in *H. pylori* N6 WT and ΔHP1021 mutant strains cultured under microaerobic and aerobic conditions (5% and 21% O2, respectively). Three independent analyses for each strain and condition are presented. **b** Comparison of HP1021-DNA interactions in the WT strain under microaerobic (red line) and aerobic (blue line) conditions with the control strain ΔHP1021 cultured under microaerobic (green line) and aerobic (purple line) conditions. The lines show mean values of reads (for 100 bp long regions) with 95% confidence intervals for 56 sites differentially bound by HP1021 protein according to edgeR analysis. For all sites, 1000 bp long fragments were extracted from the chromosome centred around the best position as determined by edgeR analysis.

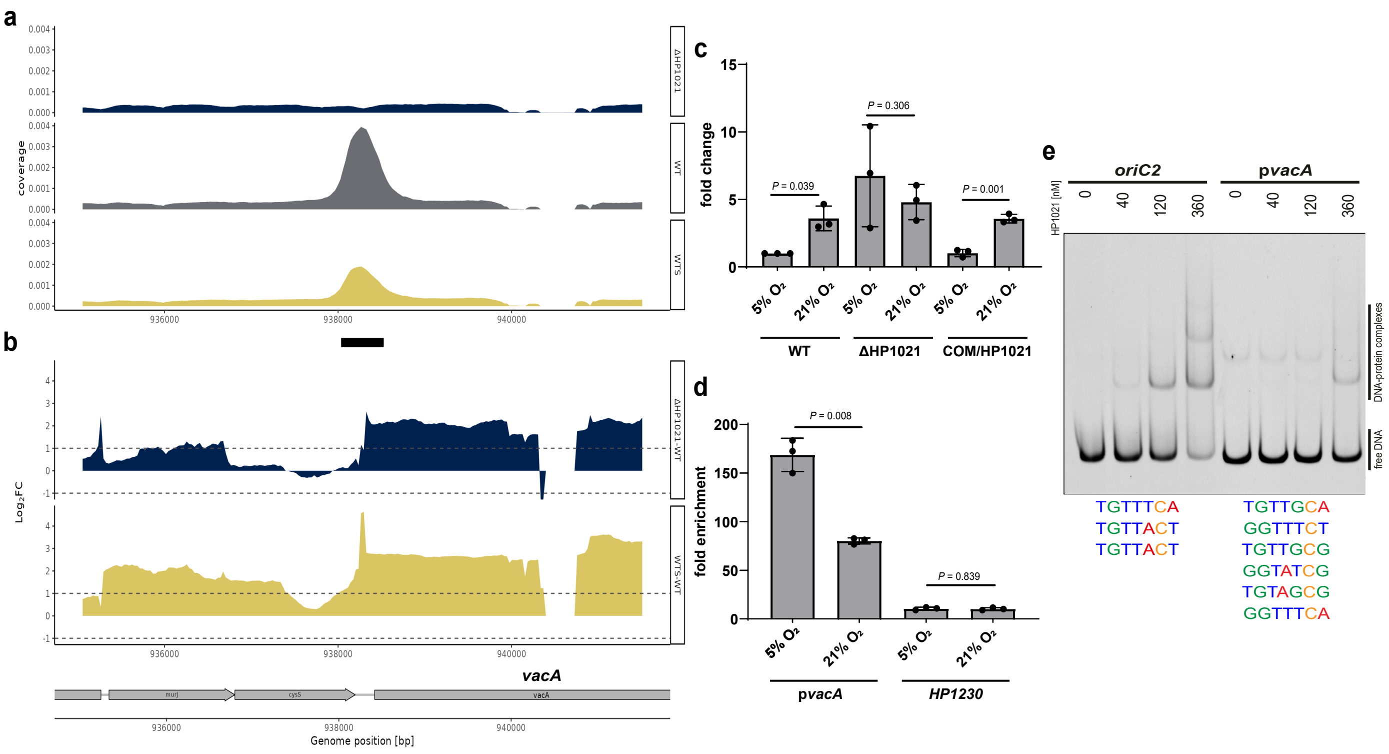


**Fig. S2: HP1021 controls *vacA* expression. a** ChIP-seq data profile of the *vacA* gene. Read counts were determined for *H. pylori* N6 WT, WTS and ΔHP1021 strains. The y-axis represents the coverage of the DNA reads, while the x-axis represents the position of the genome (in bps). The main peak of the binding site is marked with a thick black line under the x-axis. **b** RNA-seq data profile of *vacA* gene. The genomic locus for *H. pylori* N6 WT, WTS and ΔHP1021 strains with the WTS-WT and ΔHP1021-WT expression comparison; values above the black dashed lines indicate a change in the expression of $|\log_2 FC| \geq 1$; FDR $\leq 0.05$. **c** RT-qPCR analysis of the transcription of *vacA* in *H. pylori* N6 cells cultured under microaerobic and aerobic conditions (5% and 21% $O_2$, respectively). The results are presented as the fold change compared to the WT strain. **d** ChIP fold enrichment of DNA fragment in *vacA* analyzed by ChIP-qPCR in *H. pylori* N6 cells cultured under microaerobic and aerobic atmosphere (5% and 21% $O_2$, respectively). The *HP1230* gene was used as a negative control not bound by HP1021. **e** EMSA analysis of HP1021 binding to the p*vacA* region *in vitro*. EMSA was performed using the FAM-labeled DNA fragments and recombinant Strep-tagged HP1021. The *oriC2* DNA fragment was used as a high-affinity control. The HP1021 boxes (putative in the promoter p*vacA* region and experimentally determined in the *oriC2* region) are shown below the gel image. Digital processing was applied equally across the entire image, including controls. **c-d** Data are depicted as the mean values ± SD. Two-tailed Student's t-test determined the *P* value.

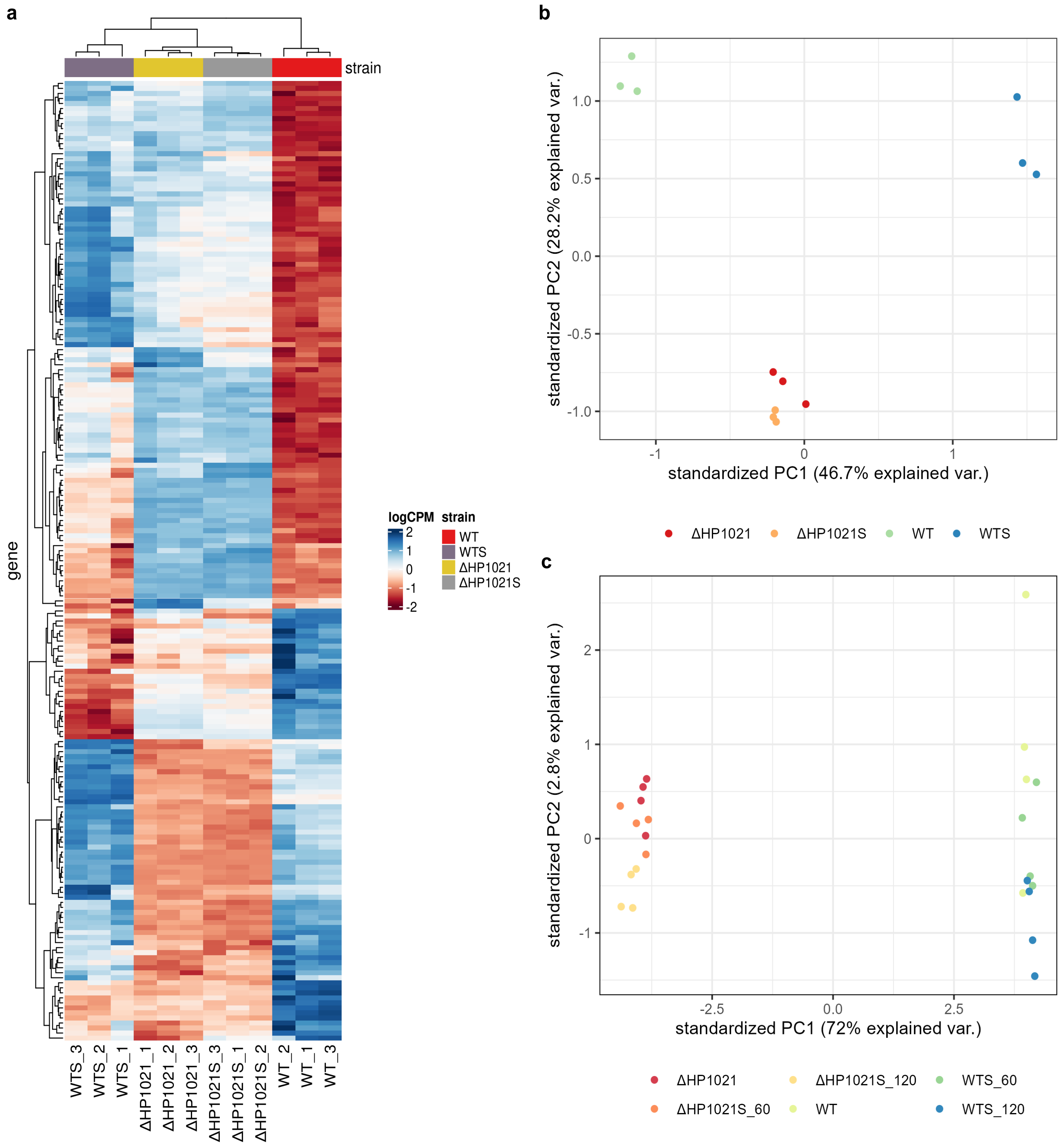


**Fig. S3: The reproducibility of biological replicates in omics data. a** Heat map of RNA-seq transcriptome. Analysis for 191 genes differentially transcribed ($|\log_2 FC| \geq 1$ and FDR $\leq 0.05$) in *H. pylori* N6 ΔHP1021 mutant strain compared to the WT strain. Plotted values are log-Counts-Per-Million (logCPM) normalized expression values of the differentially expressed gene. Data are normalized for library size and scaled to the same mean (0) and standard deviation for each gene. **b** Principal component analysis (PCA) of the normalized RNA-seq CPM data of *H. pylori* N6 strains under microaerobic and aerobic conditions (5% and 21% $O_2$, respectively). **c** PCA of the normalized proteomics CPM data of *H. pylori* N6 strains under microaerobic conditions (WT, ΔHP1021) and in response to oxidative stress after 60 minutes (WTS_60, ΔHP1021S_60 ) and 120 minutes (WTS_120, ΔHP1021S_120).

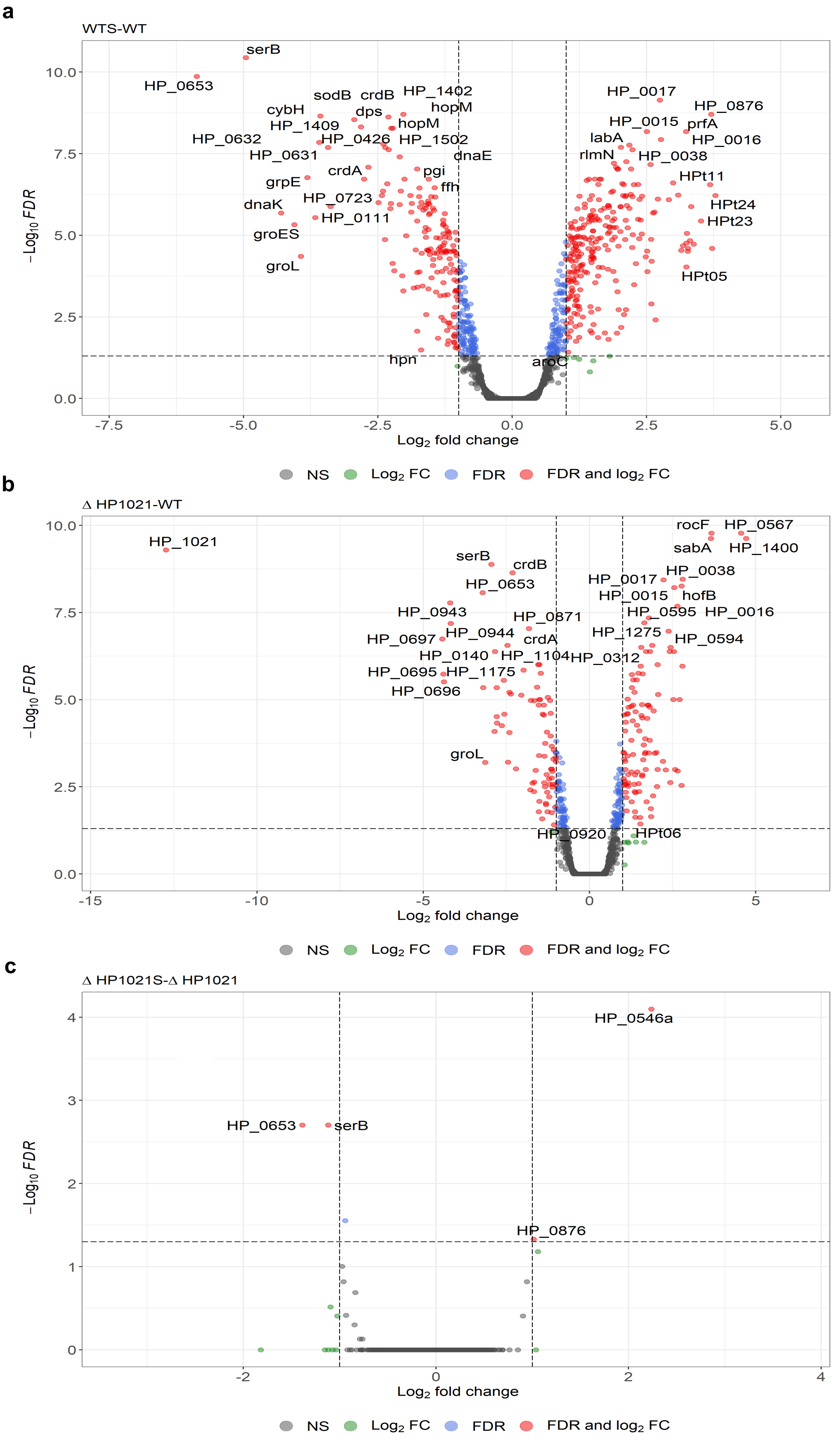


**Fig. S4: Overview of the gene regulation mediated by HP1021 revealed by RNA-seq. a** Volcano diagram of genes differently transcribed in the ΔHP1021 mutant strain compared to the wild-type (WT) strain (ΔHP1021-WT). **b** Volcano diagram of genes differently transcribed in the wild-type strain under oxidative stress (WTS) compared to the non-stressed wild-type strain (WTS-WT). **c** Volcano diagram of genes differently transcribed in the ΔHP1021 strain under oxidative stress (ΔHP1021S) compared to the non-stressed ΔHP1021 mutant (ΔHP1021S-ΔHP1021). **a-c** Three independent biological replicates were analyzed. Green dots correspond to genes with $|\log_2 FC| \geq 1$ and FDR $\geq 0.05$; blue dots correspond to genes with $|\log_2 FC| \leq 1$ and FDR $\leq 0.05$; red dots correspond to genes with $|\log_2 FC| \geq 1$ and FDR $\leq 0.05$; grey dots correspond to genes that were not significantly changed.

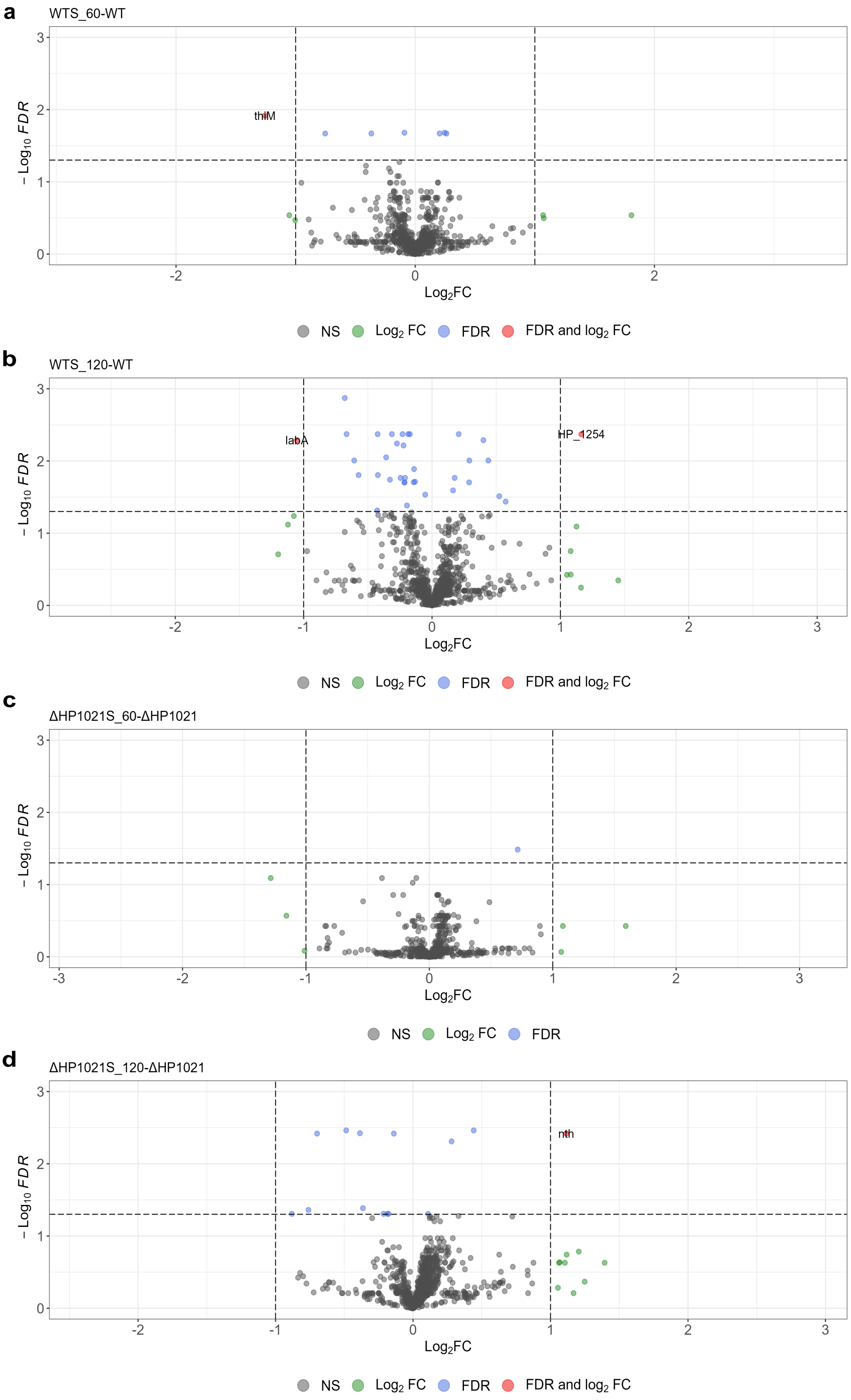


**Fig. S5: Overview of the protein level regulation mediated by HP1021 revealed by MS/LC-MS. a** Volcano diagram of proteins differentially expressed in the wild-type strain after 60-min oxidative stress (WTS_60) compared to the non-stressed WT strain (WTS_60-WT). **b** Volcano diagram of proteins differentially expressed in the wild-type strain after 120 min of oxidative stress (WTS_120) compared to the non-stressed wild-type strain (WTS_120-WT). **c** Volcano diagram of proteins differentially expressed in the ΔHP1021 mutant strain after 60 min of oxidative stress (ΔHP1021S_60) compared to the non-stressed ΔHP1021 mutant strain (ΔHP1021-ΔHP1021S_120). **d** Volcano diagram of proteins differentially expressed in the ΔHP1021 mutant strain after 120 min of oxidative stress (ΔHP1021S_120) compared to the non-stressed ΔHP1021 mutant strain (ΔHP1021S_120-ΔHP1021). **a-d** Four independent biological replicates were analyzed. Green dots correspond to genes with $|\log_2 FC| \geq 1$ and FDR $\geq 0.05$; blue dots correspond to genes with $|\log_2 FC| \leq 1$ and FDR $\leq 0.05$; red dots correspond to genes with $|\log_2 FC| \geq 1$ and FDR $\leq 0.05$; grey dots correspond to genes that were not significantly changed.

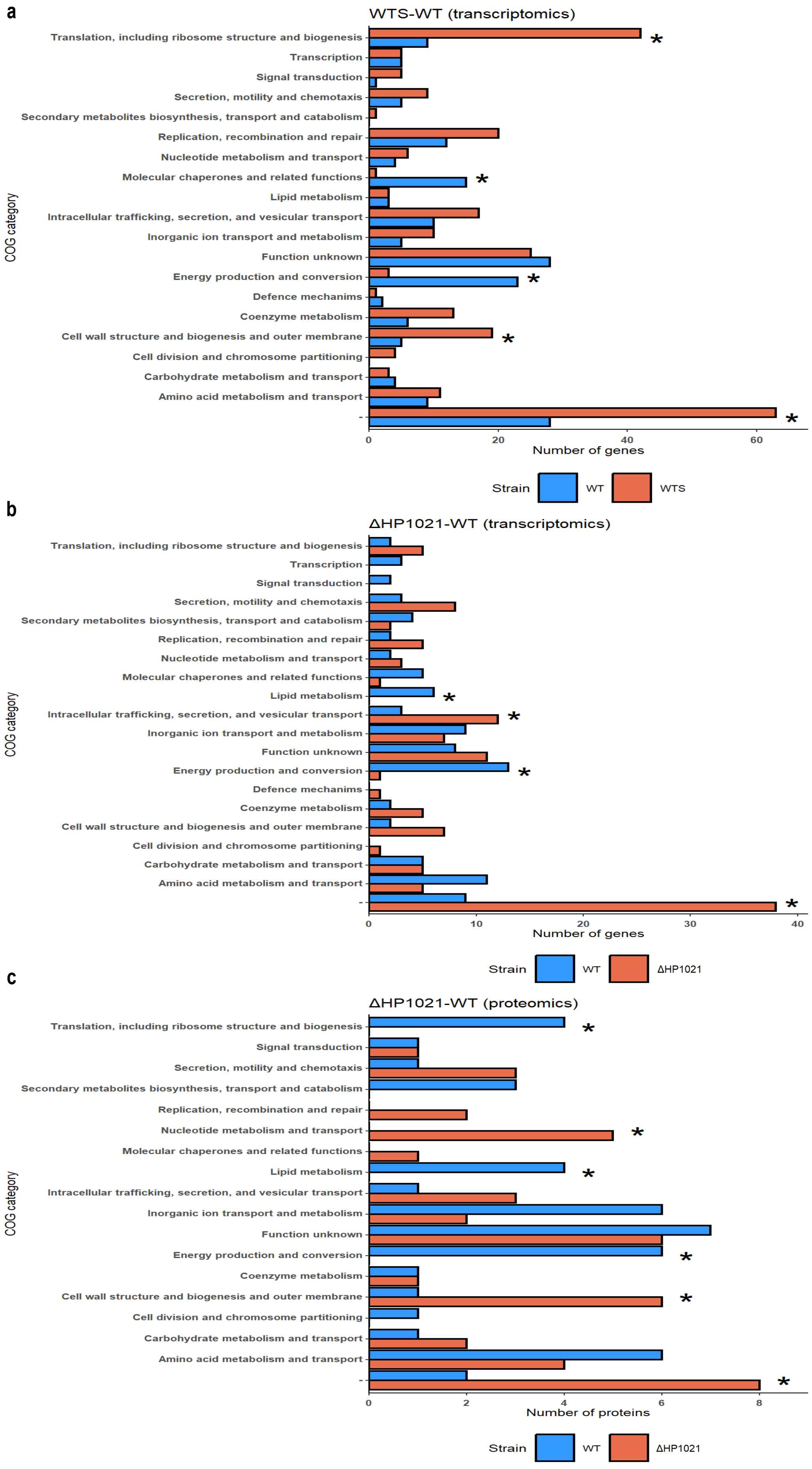


**Fig. S6:** ***H. pylori*** **N6 Clusters of Orthologous Groups (COG). a** COG groups of genes differently transcribed in the wild-type strain under oxidative stress (WTS) compared to the non-stressed wild-type strain (WTS-WT). **b** COGgroups of genes differently transcribed in the ΔHP1021 mutant strain compared to the wild-type (WT) strain (ΔHP1021-WT). **c** COGgroups of proteins expressed differently transcribed in the ΔHP1021 mutant strain compared to the wild-type (WT) strain (ΔHP1021-WT). **a-b** Chi-squared test determined the P value; significantly affected COGs ($P \leq 0.05$) are marked with black stars.

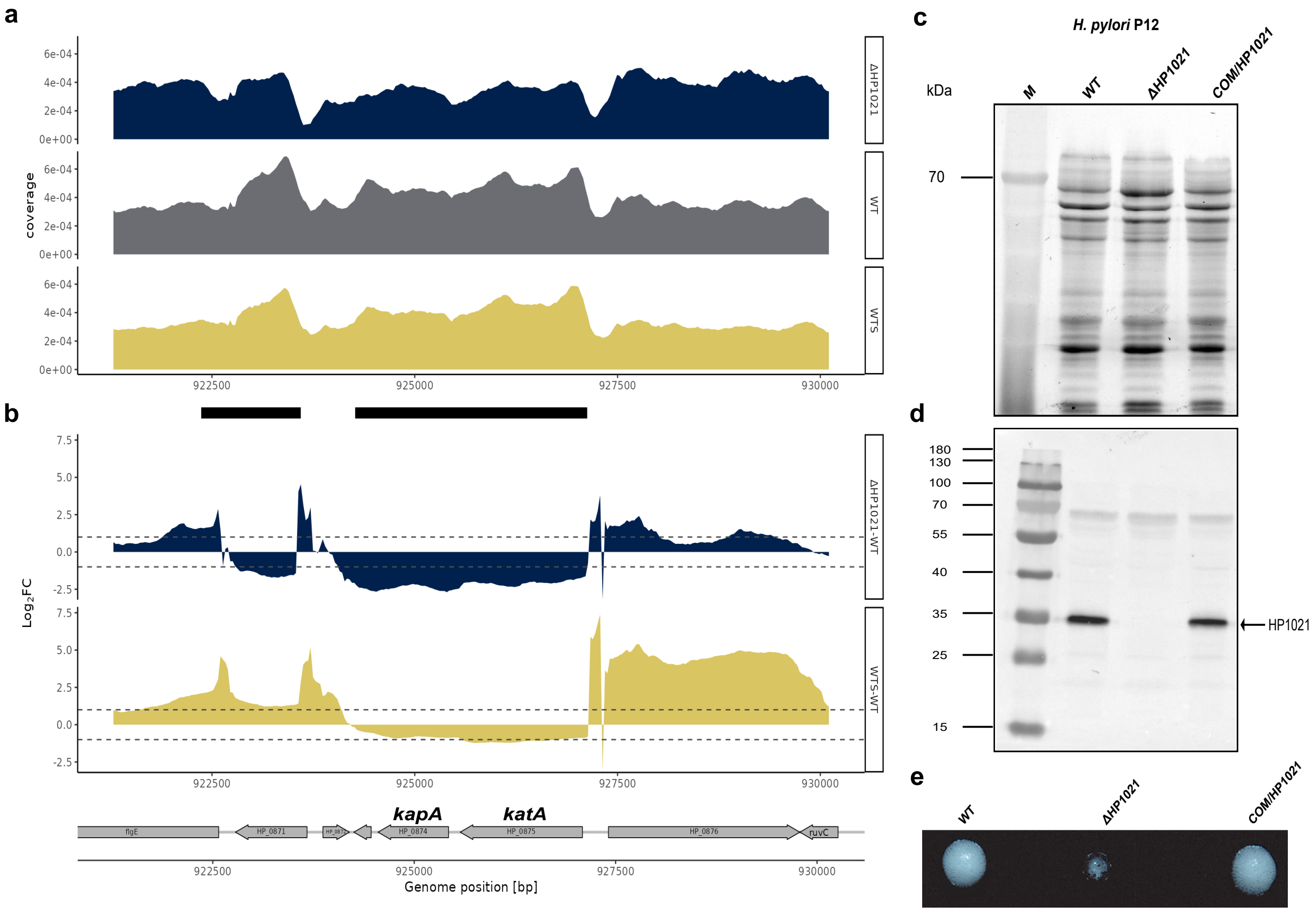


**Fig. S7: HP1021 controls *katA* and *kapA* expression. a** ChIP-seq data profile of the *katA* and *kapA* genes. Read counts were determined for *H. pylori* N6 WT, WTS and ΔHP1021 strains. The y-axis represents the coverage of the DNA reads, while the x-axis represents the position of the genome (in bps). The main peak of the binding site is marked with a thick black line under the x-axis. **b** RNA-seq data profile of *katA* and *kapA* genes. The genomic locus for *H. pylori* N6 WT, WTS and ΔHP1021 strains with the WTS-WT and ΔHP1021-WT expression comparison; values above the black dashed lines indicate a change in the expression of $|\log_2 FC| \geq 1$; FDR $\leq 0.05$. **c** Western blot analysis of HP1021 in *H. pylori* P12 wild-type and mutant strains. Lysate of each *H. pylori* strain (approximately $1.4 \times 10^8$ cells per well) was resolved in a 10% SDS-PAGE gel visualized by the TCE-UV method. **d** HP1021 was detected in bacterial lysates by a rabbit polyclonal anti-6HisHP1021 IgG. The SDS-PAGE and Western blot were performed as previously[2]. M, PageRuler Prestained Protein Ladder (Thermo Fisher Scientific). **e** Liquid cultures (10 µl) of *H. pylori* P12 of similar cell density ($OD_{600}$ ~1) were treated with an equal volume of 30% $H_2O_2$. Air bubbles produced by catalase are visible as white spots. A significant decrease in foam indicates lower catalase activity in the ΔHP1021 cells. **c-e** Digital processing was applied equally across the entire image.

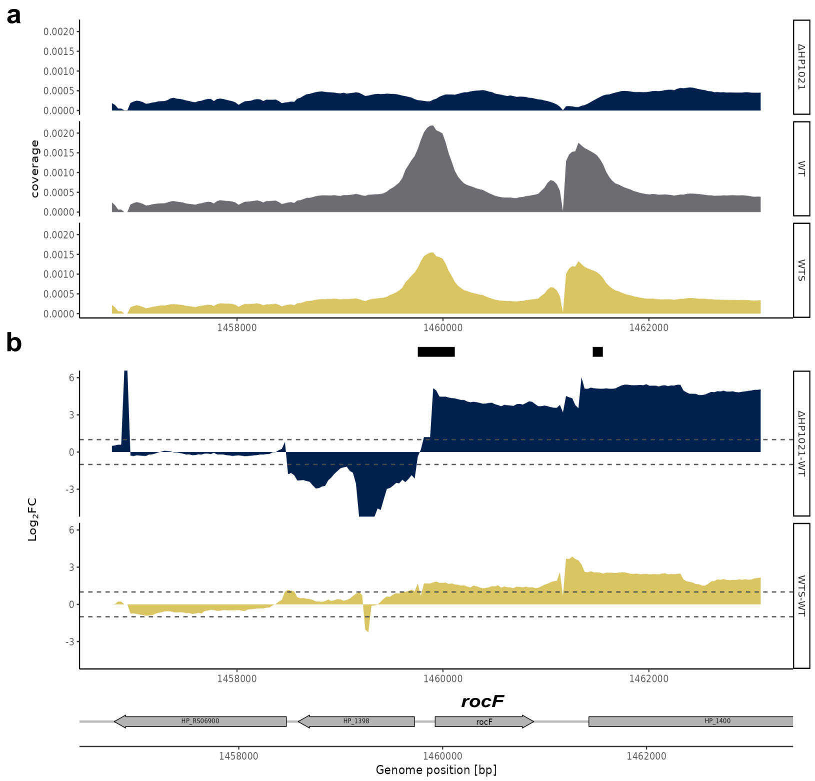


**Fig. S8: HP1021 controls *rocF* expression. a** ChIP-seq data profile of the *rocF* gene. Read counts were determined for *H. pylori* N6 WT, WTS and ΔHP1021 strains. The y-axis represents the coverage of the DNA reads, while the x-axis represents the position of the genome (in bps). The main peak of the binding site is marked with a thick black line under the x-axis. **b** RNA-seq data profile of *rocF* gene. The genomic locus for *H. pylori* N6 WT, WTS and ΔHP1021 strains with the WTS-WT and ΔHP1021-WT expression comparison; values above the black dashed lines indicate a change in the expression of $|\log_2 FC| \geq 1$; $FDR \leq 0.05$.

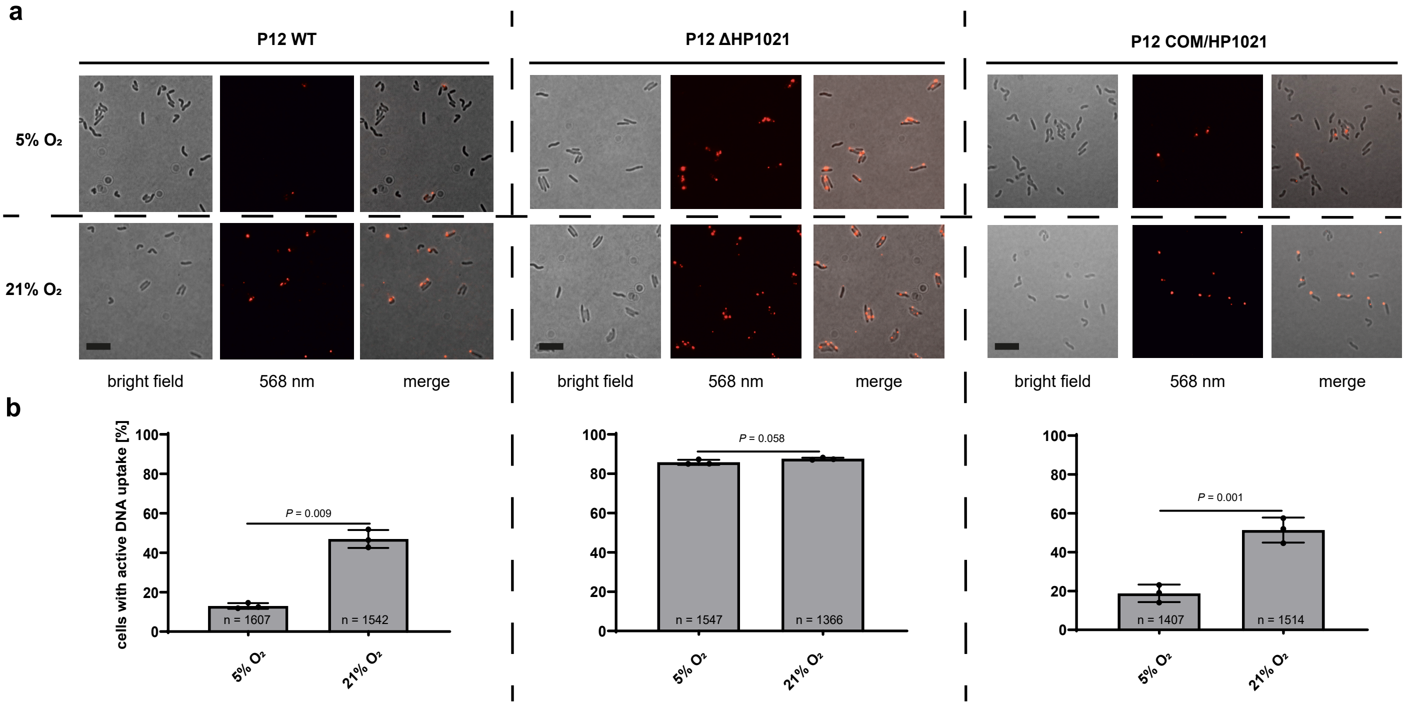


**Fig. S9: Analysis of DNA uptake by *H. pylori* P12. a** Bright field, fluorescent (532 nm) and merged images of *H. pylori* WT and mutant strains after 15 min of Cy3-λ DNA uptake under microaerobic and aerobic conditions (5% and 21% $O_2$, respectively). **b** Quantitative analysis of λ-Cy3 DNA foci formation in *H. pylori* under microaerobic and aerobic conditions (5% and 21% $O_2$, respectively). The scale bar represents 2 μm. Data are depicted as the mean values ± SD. Two-tailed Student's t-test determined the *P* value. n, sample size.

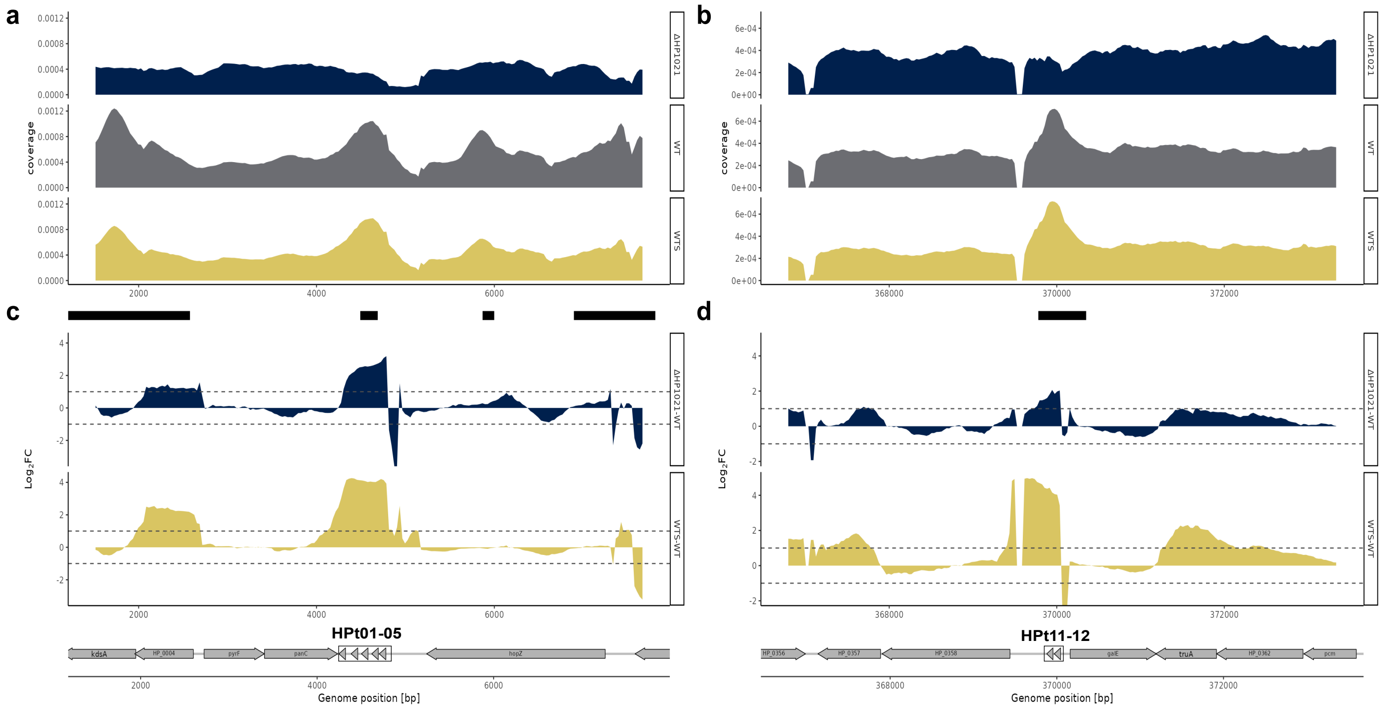


**Fig. S10: HP1021 controls tRNA expression. a-b.** ChIP-seq data profile of the regions coding tRNAs, namely (a) HPt01-HPt05 and (b) HPt11-HPt12. Read counts were determined for *H. pylori* N6 WT, WTS and ΔHP1021 strains. The y-axis represents the coverage of the DNA reads, while the x-axis represents the position of the genome (in bps). The main peak of the binding site is marked with a thick black line under the x-axis. **c-d** RNA-seq data profile of the regions coding tRNAs, namely (c) HPt01-HPt05 and (d) HPt11-HPt12. The genomic locus for *H. pylori* N6 WT, WTS and ΔHP1021 strains with the WTS-WT and ΔHP1021-WT expression comparison; values above the black dashed lines indicate a change in the expression of $|\log_2 FC| \geq 1$; FDR $\leq 0.05$.

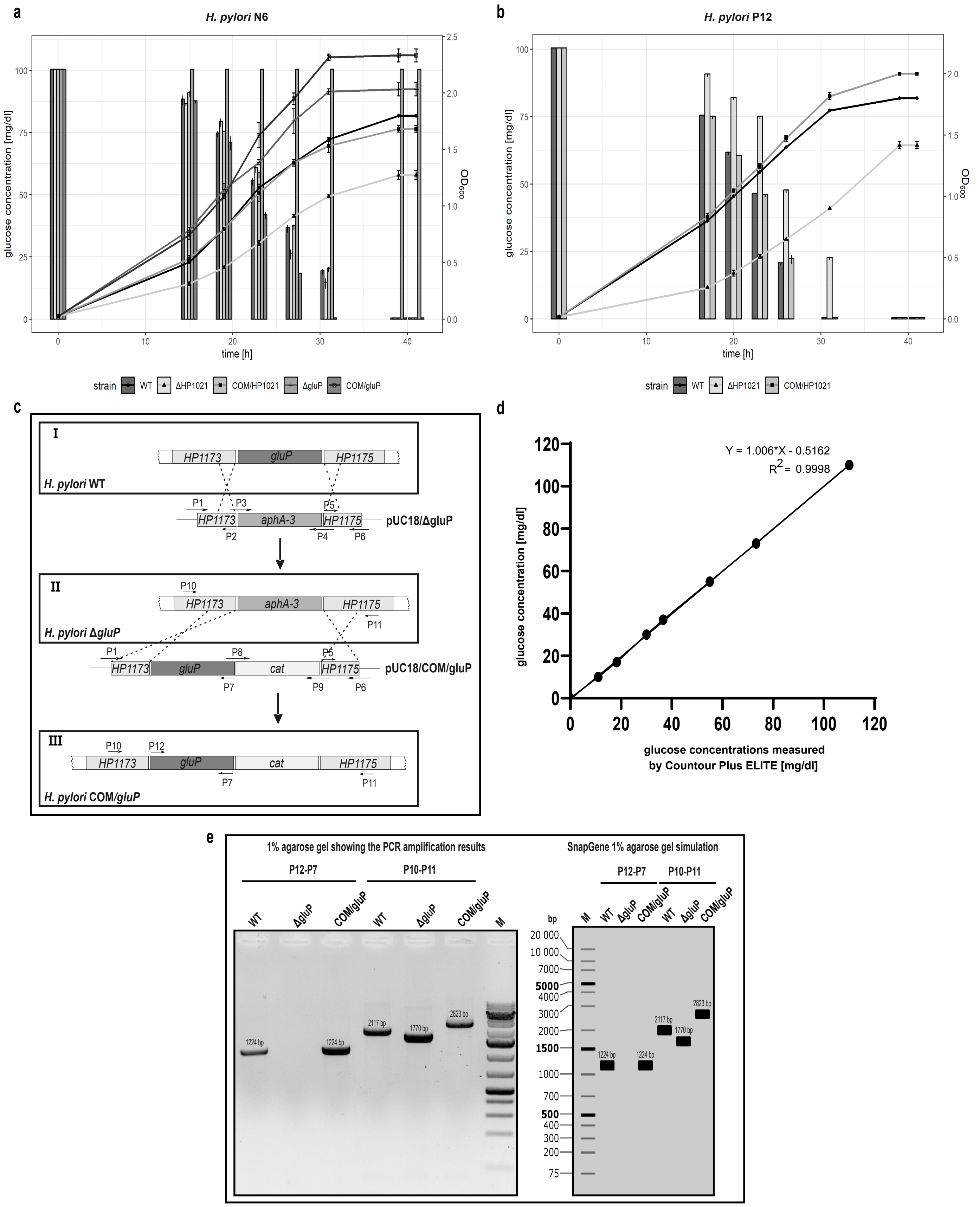


**Fig. S11: HP1021 controlled glucose uptake via GluP transporter. a** Growth curves of *H. pylori* N6 wild-type and mutant strains (line plot), combined with glucose concentration in the culture (bar plot). **b** Growth curve of *H. pylori* P12 WT, ΔHP1021, COM/HP1021 combined with glucose consumption (bar plot). **c** The mutagenesis strategy used to delete and subsequently complement *gluP* on the *H. pylori* N6 chromosome. *H. pylori* N6 wild-type *gluP* chromosomal loci (I) and plasmid DNA for double crossing-over to give *H. pylori* N6 ΔgluP (II) and COM/gluP (III) mutant strains are shown. For the plasmids and primer sequences, see Supplementary Tables S4 and S5, respectively. **d** Glucose concentration standard curve. The TSBΔD-FBS medium supplemented with glucose (110 mg/dl) was serially diluted with 1 × PBS, and the glucose concentration of the appropriate dilutions was measured with Contour Plus ELITE. **e** Agarose gel electrophoresis of PCR products confirming the correct *H. pylori* N6 ΔgluP and COM/gluP mutant strain construction with SnapGene® agarose gel simulation. M, GeneRuler™ 1 kb Plus DNA Ladder (Thermo Fisher Scientific).

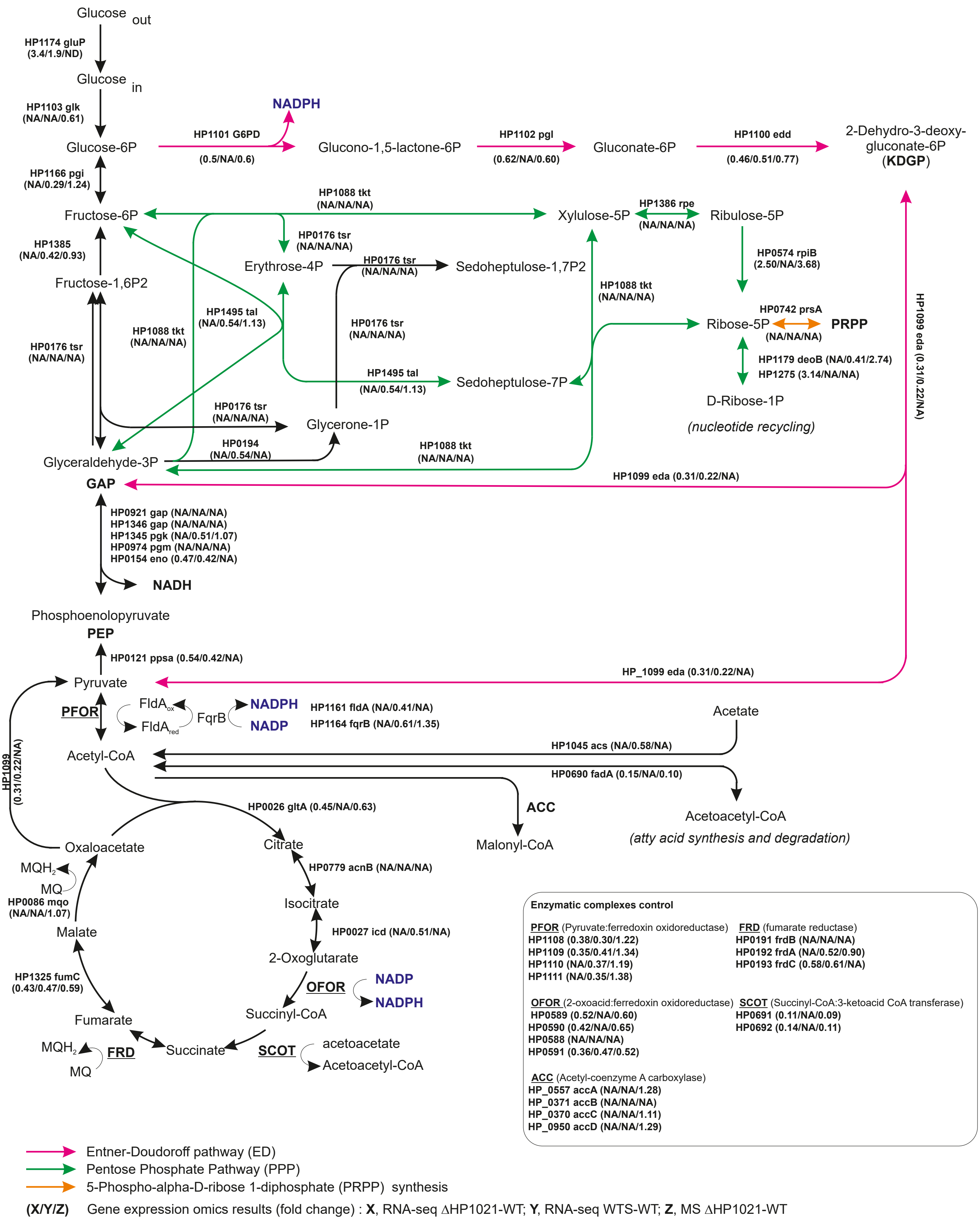


**Fig. S12: A model of glucose metabolism in *H. pylori* N6 based on KEGG database and Steiner et al.**[9]**.** Genes annotation according to *H. pylori* 26695 (NC_000915.1) strain . MQ, menaquinone; Fd, ferredoxin; NA, indifferent gene (the change was not significant); ND, not detected.

**Supplementary References**


1. Ge, Z. & Taylor, D. E. *H. pylori* DNA Transformation by Natural Competence and Electroporation. *Helicobacter pylori Protoc.* 145–152 (1997).
2. Szczepanowski, P. *et al.* HP1021 is a redox switch protein identified in *Helicobacter pylori*. *Nucleic Acids Res.* 49, 6863–6879 (2021).
3. Donczew, R. *et al.* The atypical response regulator HP1021 controls formation of the *Helicobacter pylori* replication initiation complex. *Mol. Microbiol.* 95, 297–312 (2015).
4. Gibson, D. G. *et al.* Enzymatic assembly of DNA molecules up to several hundred kilobases. *Nat. Methods* 65 6, 343–345 (2009).
5. Sambrook, J. & Russel, D. W. Molecular Cloning: A Laboratory Manual. *Cold Spring Harbor Laboratory Press* (2001).
6. Tomb, J. F. *et al.* The complete genome sequence of the gastric pathogen *Helicobacter pylori*. *Nature* 388, 539–547 (1997).
7. Ferrero, R. L., Cussac, V., Courcoux, P. & Labigne, A. Construction of isogenic urease-negative mutants of *Helicobacter pylori* by allelic exchange. *J. Bacteriol.* 174, 4212–4217 (1992).
8. Donczew, R., Weigel, C., Lurz, R., Zakrzewska-Czerwińska, J. & Zawilak-Pawlik, A. *Helicobacter pylori oriC* - the first bipartite origin of chromosome replication in Gram-negative bacteria. *Nucleic Acids Res.* 40, 9647 (2012).
9. Steiner, T. M. *et al.* Substrate usage determines carbon flux via the citrate cycle in *Helicobacter pylori*. *Mol. Microbiol.* 116, 841–860 (2021).
